# *In vivo* experimental model of β-glucan-induced innate immune memory

**DOI:** 10.64898/2026.02.13.705736

**Authors:** Khrystyna Matvieieva, Sarah Sze Wah Wong, Mathieu Bohm, Ali Hassan, Jessica Quintin

## Abstract

**SUMMARY:** Innate immune memory is the ability of innate immune cells to develop a recallable, epigenetically imprinted response after an initial stimulus, enabling them to mount differential responses upon restimulation. Here, we present a detailed protocol of β-glucan preparation, and its *in vivo* application to modulate innate immune memory in mice through repeated intraperitoneal (IP) injections. This approach results in enhanced hematopoiesis and bone marrow specific myeloid bias, key hallmarks of *in vivo* innate immune memory responses.

For complete details on the use and execution of this protocol, please refer to Hassan et al.

**Graphical abstract:** 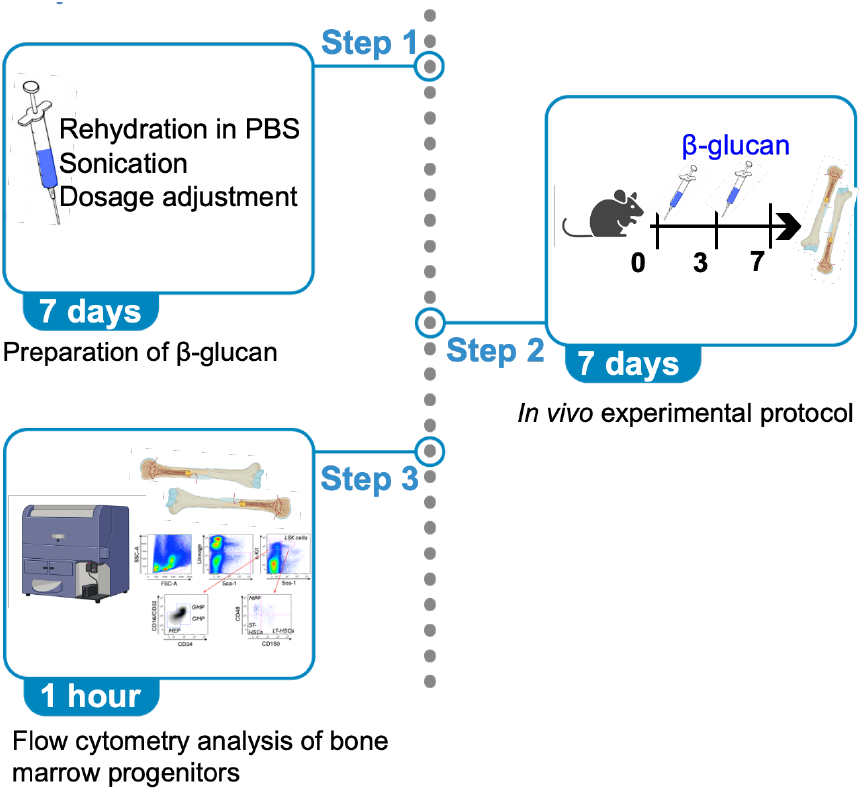

**Highlights:**

- Preparation of β-glucan for *in vivo* experiments.
- *In vivo* modulation of innate immune memory through repeated intraperitoneal β-glucan injections in mice.
- Modulation of innate immune memory is assessed by enhanced bone marrow hematopoiesis and GMP production.

## BEFORE YOU BEGIN

The capacity of innate immune cells to acquire memory-like properties was first described more than a decade ago and is known as innate immune memory. This phenomenon reflects the ability of innate immune cells to undergo epigenetic and metabolic reprogramming after an initial stimulus, enabling them to mount differential responses upon subsequent challenges, either through trained immunity (enhanced responsiveness) or tolerance (reduced responsiveness)^1,2,3^. β-glucan, a polysaccharide abundant in the cell walls of microbes and plants, is among the most extensively studied inducers of innate immune memory.

Several *in vitro* studies have explored the mechanisms underlying trained immunity, particularly using adherent human monocytes stimulated with β-glucan derived from *Candida albicans*^4,5,6^. Detailed and standardized protocol for the *in vitro* induction of trained immunity is available^7^, whereas standardized protocols for *in vivo* mouse models are still lacking. Nevertheless, studies using different β-glucan preparations have become increasingly common over the past few years. In the last five years alone, over 500 *in vivo* studies have tested β-glucans in diverse contexts, including tumor and infection models, metabolic and dietary interventions, vaccine adjuvants, and innate immune memory/trained immunity research. Among the studies focusing on β-glucan-induced innate immune memory/trained immunity, different β-glucan preparations and sources were used^8,9,10,11,12^. Notably, depending on the source of β-glucan, structural properties may differ, including particle size, molecular weight, purity, and branching characteristics (such as type, length, and frequency of branches), which can in turn impact the biological activity of β-glucan. Thus, a standardized *in vivo* protocol that includes all aspects of β-glucan preparation and administration, along with structural characterization, is needed.

In this study, we used β-glucan from Immune Health Basics (Wellmune), a 1,3/1,6-β-glucan derived from *Saccharomyces cerevisiae*.

## Institutional permissions

All procedures involving mice, including breeding and experiments, were conducted in accordance with the guidelines of the Institut Pasteur Ethics Committee and were approved by the French Ministry of Research under ethical protocols CETEA: 180090 – 240035.

## KEY RESOURCES TABLE

| REAGENT or RESOURCE | SOURCE | IDENTIFIER |
| --- | --- | --- |
| **Antibodies** |  |  |
| Lineage: |  |  |
| CD3 (clone 145-2C11) | Biolegend | Cat#100308; RRID: AB_312673 |
| CD19 (clone 1D3) | Invitrogen | Cat# 12-0193-82; RRID: AB_657659 |
| NK1.1 (clone 29A1.4) | Invitrogen | Cat#12-3351-82; RRID: AB_1210743 |
| Ly6G (clone 1A8) | BD Biosciences | Cat# 551459; RRID: AB_394206 |
| GR-1 (clone RB6-8C5) | Biolegend | Cat#108408; RRID: AB_313373 |
| TER-119 (clone TER-119) | Biolegend | Cat#116202; RRID: AB_313709 |
| CD11b (clone M1/70) | BD Biosciences | Cat# 557397; RRID: AB_396680 |
| Surface markers: |  |  |
| Flt3 (clone A2E10.1) | BD Biosciences | Cat# 562537; RRID: AB_2737639 |
| Sca-1 (clone D7) | BD Biosciences | Cat# 562729; RRID: AB_2737750 |
| c-kit (clone 2B8) | BD Biosciences | Cat#563146; RRID: AB_2738028 |
| CD48 (clone HM48-1) | Biolegend | Cat#103412; RRID: AB_571997 |
| CD150 (clone TC15-12F12.2) | Biolegend | Cat# 115914; RRID: AB_439797 |
| CD34 (clone RAM34) | Invitrogen | Cat#56-0341-82; RRID: AB_493998 |
| CD16/32 (clone 93) | Biolegend | Cat#101324; RRID: AB_1877267 |
| **Biological samples** |  |  |
| Fetal bovine serum (FBS) | Sigma-Aldrich | F7524 |
| Bovine serum albumin (BSA) | Sigma-Aldrich | A7030-100G |
| **Chemicals, peptides, and recombinant proteins** |  |  |
| Phosphate buffered saline (PBS) (1x) | Gibco | Cat#10010023 |
| RPMI 1640 medium | Gibco | Cat# 11875093 |
| Ethylenediaminetetraacetic acid (EDTA) 0.5M sterile solution | Bio Basic Inc. | SD8135 |
| Formaldehyde solution | Thermo Scientific | Cat#28908 |
| β-1,3/1,6-glucan derived from *S. cerevisiae* | Immune Health Basics | J-23-5; 0149D9 |
| Critical commercial assays |  |  |
| Flow Cytometry Size Calibration Kit (nonfluorescent microspheres) | Invitrogen | Cat#F13838 |
| AccuCheck Counting Beads | Life Technologies | Cat#50-113-7556 |
| Experimental models: Organisms/strains |  |  |
| Mouse: *Balb/cByJ* | Charles River Laboratory | Female, 8-week-old |
| Software and algorithms |  |  |
| Fiji ImageJ2 (version 2.9.0/1.53t) |  | RRID:SCR_002285<br>https://imagej.net/downloads |
| FlowJo (Version 10.10.0) | Becton Dickinson | RRID: [SCR_008520](https://www.flowjo.com/)<br>https://www.flowjo.com/ |
| Prism (Version 10.4.1) | GraphPad Software Inc. | RRID:SCR_005375 |

### Materials and equipment

#### FACS buffer

Materials and equipment FACS buffer
| Reagent | Final concentration | Amount |
| --- | --- | --- |
| BSA | 0.5% | 2.5 mL |
| EDTA (0.5M) | 2mM | 2 mL |
| PBS | n/a | 495.5 mL |
| **Total** | **n/a** | **500 mL** |

Store at 4°C for up to 3 months

## STEP-BY-STEP METHOD DETAILS

### Step 1: Preparation of β-glucan suspension

#### Timing

7 days

#### Before starting

Open the Wellmune β-glucan capsule under a biosafety hood using sterile forceps and transfer powder into a 50 mL conical tube. After use, seal the tube with parafilm and store it at 4 °C for up to three months.

##### 1. Day 0

Weighing of β-glucan powder

a. Under the biosafety hood, disinfect the spatula by placing it in a 50 mL conical tube filled with ANIOS or another disinfectant. Leave it immersed for 5–10 minutes.
b. Transfer the estimated amount of β-glucan powder into a 15 mL tube and weigh it.
c. Label the tube with the amount in mg and add 1 mL of sterile PBS (under the hood).
d. Store the suspension at 4 °C for 5 to 7 days.

#### Note

Prolonged incubation of β-glucan in PBS is essential for rehydrating the powder and restoring its native conformation in aqueous environments, thereby enhancing its suitability for biological and immunological applications. Rehydration also reduces aggregation. Consistently, we observed a visibly more homogenous and “fluffier” suspension following PBS incubation (Figure 1A).

**Figure 1.**
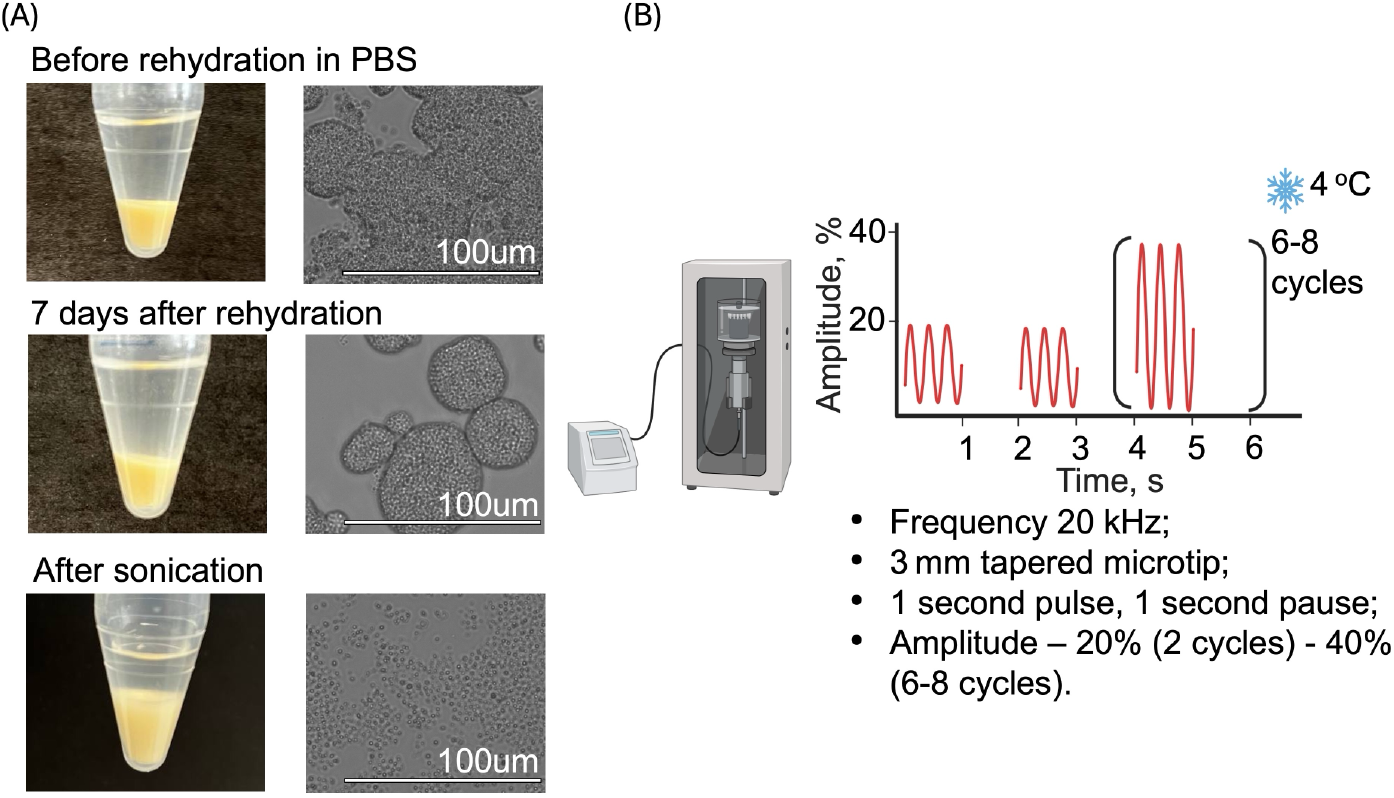
β-glucan preparation. Macroscopic (left) and microscopic images (right) of β-glucan at different steps of preparation. Detailed sonication protocol.

##### 2. Day 7: Sonication

Sonication is a critical processing step that reduces both the particle size and molecular weight of β-glucan, key parameters for safe and effective *in vivo* use. In contrast, administration of non-sonicated, particulate β-glucan (e.g., zymosan) has been associated with adverse effects, such as hyperinflammation, pain, and stress responses in mice^13^.

a. Under a biosafety hood, prepare four 50 mL conical tubes as follows: 50 mL of ANIOS or another disinfectant; 50 mL of 70% Ethanol; Two tubes containing 50 mL of sterile PBS, labeled PBS 1 and PBS 2.
b. Disinfect the sonicator probe by immersing it in disinfectant for a minimum of 5 minutes.
c. Sequentially rinse the probe by dipping it into the 70% Ethanol tube, followed by PBS 1, then PBS 2. Gently remove any residual PBS from the probe tip using sterile paper.
d. Set the sonicator to a 20 kHz frequency using a 3 mm tapered microtip and apply 1-second pulses with 1-second pauses.
e. Insert the probe to the 15 mL tube containing β-glucan powder and 1 mL of sterile PBS.
f. Begin sonication at 20% amplitude, then increase to 40% after two cycles and continue for a total of 8–10 cycles, as illustrated in Figure 1B.

#### Note

Ensure that the sonicator probe is inserted into the solution. The volume of sterile PBS (1 mL) is optimal to leave sufficient headspace to prevent overflowing, and thus sample loss, during sonication.

#### Note

Alternative sonication protocol (e.g., 45-second pulses with 15-second pauses at 20% amplitude for 12 cycles) is also effective.

#### Note

Multiple sonicator systems - Vibra-Cell, B-30, and Branson 250 - were tested, all yielding comparable results.

#### Note

The sonicated β-glucan solution exhibited improved dispersibility and a marked reduction in particle size and aggregation (Figure 1A).

g. Immediately reseal the tube and place it back on ice or store at 4 °C.
h. Clean the probe using the same tube sequence as in step c: disinfectant, 70% ethanol, PBS 1, PBS 2. Dry the probe thoroughly with sterile paper.

#### Optional

Measurement of β-glucan particles size (see Table 1)

**Table 1.** Size of β-glucan particles measured using different methods.

| Condition |  | Method |  |  |  |  |  |
| --- | --- | --- | --- | --- | --- | --- | --- |
|  |  | Optical microscopy + Fiji analysis |  |  | Flow Cytometry |  |  |
|  |  | Min. | Max. | Average | Min. | Max. | Average |
| β-glucan before rehydration with PBS | | Accurate measurement was not possible (big aggregations) | | | Accurate measurement was not possible (big particles) | | |
| β-glucan 7 days after rehydration with PBS | Before sonication | 110 | 5500 | 1400 | | | |
|  | After sonication | 0.8 | 7 | 1.4 | 1 | 5 | 1.8 |

#### Fiji software

Using the particle analysis method in Fiji, the average particle size prior to sonication was 1400 µm (range: 110–5500 µm). After sonication, the mean size decreased significantly to approximately 1.4 µm, and the suspension appeared considerably more homogeneous.

#### Flow cytometry

Using a size calibration kit (according to the manufacturer’s instructions), most post-sonication particles were 2 µm in diameter (55%), followed by 1 µm (29%), 3 µm (10%), 4 µm (1%), and >4 µm (0.9%). These results confirm that our protocol produces a relatively uniform particle suspension suitable for *in vivo* experimentation.

#### Optional

Assessment of β-glucan structure (see Table 2)

**Table 2.**
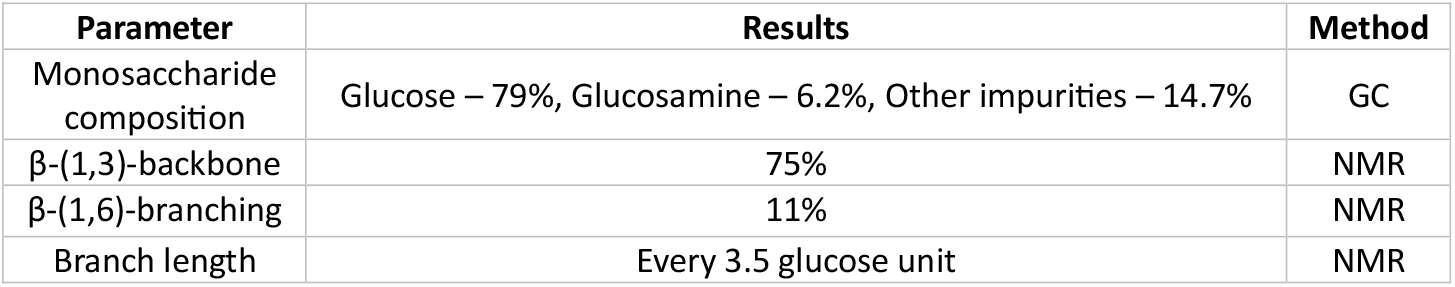
Example of β-glucan structural assessment by gas chromatography (GC) and nuclear magnetic resonance (NMR)

| Parameter | Results | Method |
| --- | --- | --- |
| Monosaccharide composition | Glucose – 79%, Glucosamine – 6.2%, Other impurities – 14.7% | GC |
| β-(1,3)-backbone | 75% | NMR |
| β-(1,6)-branching | 11% | NMR |
| Branch length | Every 3.5 glucose unit | NMR |

#### Gas chromatography (GC)

Using established methods described in previous studies^14,15^, GC can be employed to evaluate the monosaccharide composition of β-glucan. Pure β-glucan preparations typically contain glucose as the dominant monosaccharide. Despite the detection of minor lipid and glucosamine residues, our β-glucan preparation is regarded as pure, consisting predominantly of glucose units.

#### Nuclear magnetic resonance (NMR) analysis

According to the guidelines of our local analytical platform and following previously established protocols^16, 17^, the branching properties of β-glucan can be assessed by NMR spectroscopy, with samples dissolved in deuterated DMSO supplemented with 8% D_2_O. The β-glucan from our study consists predominantly of β-(1,3)-glycosidic linkages forming the main backbone, with β-(1,6)-linked side branches. On average, branches occur every ∼11 residues (11% branching) with an average branch length of approximately 3.5 glucose units. Moreover, NMR analyses performed before and after sonication confirm that sonication does not alter the branching properties of β-glucan.

#### Note

Given the strong influence of β-glucan structure on its biological activity, we strongly recommend obtaining and documenting the characteristics of your β-glucan in a similar way before starting the experiment

### Step 2: In vivo experimental protocol

#### Timing

7 days

##### 1. Day 0

First IP injection (right after the sonication step)

a. Dilute the β-glucan suspension under sterile conditions to a final concentration of 5 mg/mL using sterile PBS.
b. Mix thoroughly and load 200 µL (1 mg) of suspension into each using U-100 insulin syringe with 30G needle (one per mouse).
c. Prepare PBS-only syringes similarly for control mice.
d. Inject mice intraperitoneally.
e. Store remaining β-glucan suspension at 4°C.

#### Note

In our study, we used female Balb/cByJ mice at 8 weeks of age with ∼20 g weight in line with similar published studies, allowing for historical data comparison if needed. Indeed, Balb/cByJ strain is widely used in immunological research due to its well-characterized immune profile and better compatibility in group housing.

#### Note

To avoid the flawed variation of the Cage-Confounded Design that occurs when each treatment group is assigned to a single cage of animals, we always house both β-glucan-treated and control mice together within the same cage. If treatment group is assigned to a single cage, any observed differences among the outcomes of interest may stem from either treatment effects, cage effects or some combination of the two. Therefore, the variance attributable to treatment cannot be isolated from that attributable to the cage environment.

##### 2. Day 3

Second IP injection

a. Repeat steps 1b-d as described above.

##### 3. Day 7

Bone marrow collection, single cell suspension preparation and staining

a. Euthanize mice using cervical dislocation, CO_2_ asphyxiation, or another approved method.
b. Collect femurs and tibias from both hind limbs.
c. Cut off both ends of each bone to access the bone marrow.
d. Flush the bone marrow with a total of 10 mL RPMI 1640 medium supplemented with 10% FBS using a 5 mL syringe and 26G needle (5 mL per leg).
e. Centrifuge at 300 × g for 7 min at 4 °C.
f. Discard the supernatant.
g. Resuspend the pellet in 1 mL FACS buffer and filter through a 70 µm cell strainer.
h. Seed 50 µL of the suspension into each well of a 96-well U-bottom plate.
i. Centrifuge at 400 × g for 2 min at 4 °C.
j. Remove the supernatant by quickly inverting the plate and gently tapping it on absorbent paper.
k. Stain cells with 30 µL of antibody mix (lineage cocktail and surface markers) for 30 min at 4 °C.
l. Wash with 100 µL FACS buffer.
m. Centrifuge at 400 × g for 2 min at 4 °C.
n. Remove the supernatant as described in step j.
o. Resuspend in 100 µL FACS buffer.
p. Analyse samples on the same day following the gating strategy shown in Figure 2B. If immediate analysis is not possible, fix cells by adding 100 µL of 4% formaldehyde solution.

**Figure 2.**
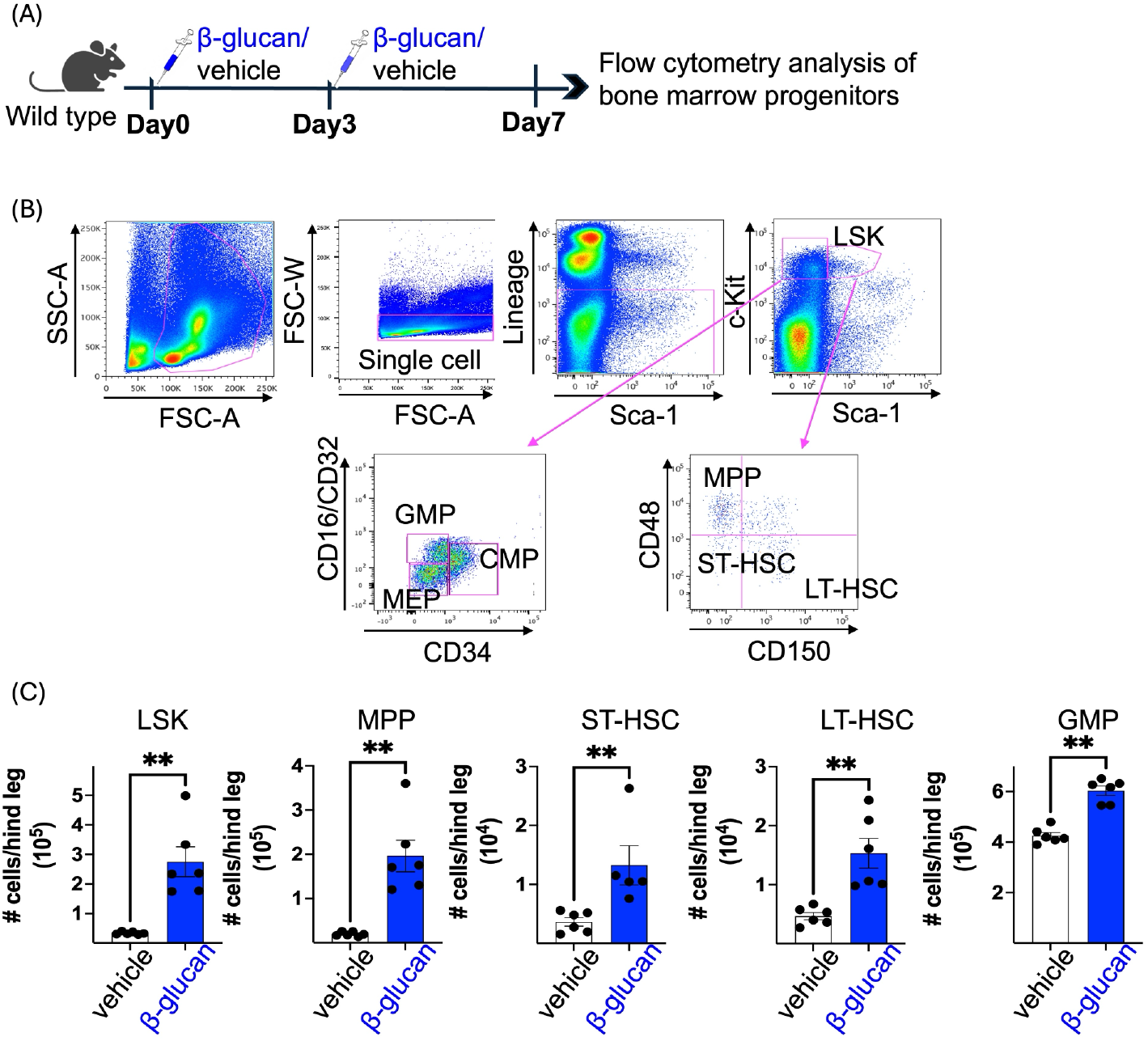
β-glucan induces innate immune memory in mice. *(A)In vivo* experimental design. Mice were injected intraperitoneally with PBS or β-glucan at day 0 and day 3. At day 7, the number of different bone marrow progenitors were determined by flow cytometry. (B)Gating strategy. Cells were gated for SSC-A against FSC-A. Mature cells and erythrocytes were excluded by selecting the lineage-negative population. Subsequent gating steps were performed according to the expression of specific markers defining different bone marrow progenitor populations. (C)Total number of different bone marrow progenitors at day 7 after β-glucan injection (n=6). Data are presented as mean ± SEM. **p<0.01; unpaired two-tailed Mann-Whitney test. LSK - Lin^−^Sca-1^+^c-Kit^+^ cells; MPP - multipotent progenitors, ST-HSC - short-term hematopoietic stem cells, LT-HSC - long-term hematopoietic stem cells, GMP - granulocyte-monocyte progenitor.

#### Critical

Samples should be analysed within 3 days of fixation to prevent degradation of dye tandems, maintain dye stability, and preserve cell viability.

### Step 3. Assessment of in vivo innate memory modulation by flow cytometry analysis of bone marrow progenitors

#### Timing

1 hour

#### Flow cytometry analysis

All flow cytometry analyses were performed on a BD LSR Fortessa (BD Biosciences), and data were analyzed using FlowJo v10 software (FlowJo LLC). Bone marrow progenitor populations were identified using a well-established gating strategy (Figure 2B)^8^.

#### Cell counting

Cell numbers for each sample were obtained using AccuCheck Counting Beads (Life Technologies), following the manufacturer’s instructions.

## EXPECTED OUTCOMES

As evidence of innate memory induction, we observed significantly enhanced hematopoiesis, characterized by increased numbers of LSK (Lin^−^Sca-1^+^c-Kit^+^), MPP (multipotent progenitors), ST-HSC (short-term hematopoietic stem cells), LT-HSC (long-term hematopoietic stem cells), and GMP (granulocyte-monocyte progenitor) (Figure 2C).

## QUANTIFICATION AND STATISTICAL ANALYSIS

All data are presented as mean ± SEM. Statistical evaluation of differences between the experimental groups was done using Mann-Whitney unpaired two-tailed U test and performed using Prism v9 (GraphPad Software Inc.). Significance was set at p < 0.05.

## LIMITATIONS

This protocol provides a detailed approach for inducing innate immune memory in mice. However, several limitations should be noted.

As a hallmark of innate immune memory, we assessed enhanced hematopoiesis and myelopoiesis in the bone marrow. Other relevant readouts of innate immune memory modulation can also be employed.

This protocol was optimized using female Balb/cByJ mice aged 8–12 weeks allowing for historical data comparison. Other variables such as mouse strain, microbiota, sex, and age have yet to be explored to obtain an exhaustive and possibly universal protocol applicable in clinics.

In this study, β-glucan was administered via IP injection. Other routes, such as intravenous or subcutaneous administration, which may be more relevant for potential human applications, were not evaluated and warrant further investigation. Additionally, we used two IP injections, whereas some published studies rely on a single IP injection ^8^,^9^ or three and more ^12^. A direct comparison between single and multiple doses would be of significant interest.

## TROUBLESHOOTING

### Problem 1

β-glucan particles do not reduce to the desired size of 1–2 µm, with visible aggregation remaining.

#### Potential solution

Repeat the sonication step using optimized settings for amplitude, cycle duration, and total time. Ensure the sonicator is functioning properly and that the probe is correctly immersed in the suspension down to the solubilized powder itself.

### Problem 2

Mice experience weight loss or death following IP injection.

#### Potential solution

This is a rare occurrence and has not been directly associated with β-glucan administration alone. Potential causes include:

1. Contamination of the β-glucan suspension – Ensure that all preparation steps are performed under sterile conditions. The sonicator probe must be thoroughly disinfected before and after use.
2. Inadequate sonication – Poor sonication may leave β-glucan aggregates that can cause adverse effects. Carefully follow the sonication protocol and verify the absence of aggregates using a microscope.
3. Incorrect IP injection technique.

Mice should be closely monitored for critical endpoints daily, including changes in grooming behavior, whisker position, posture, facial expression, signs of dehydration (e.g., skin tenting), and weight loss. A loss of more than 20% of body weight is considered a humane endpoint and requires immediate intervention.

### Problem 3

Lack of induction of innate immune memory

#### Potential solution

This issue often stems from the same causes listed in Problem 2. Ensure correct β-glucan preparation and administration. If issues persist, repeat the experiment with strict adherence to the protocol, especially the sonication and dosing steps.

### Problem 4

High variability in bone marrow cell counts between mice.

#### Potential solution

Some variability is natural due to individual differences among mice. However, inaccuracies may also result from inconsistent use of counting beads. To minimize this, ensure that beads are well mixed before use, added in consistent volumes across samples, and that acquisition settings remain uniform during analysis.

## RESOURCE AVAILABILITY

### Lead contact

Further information and requests for resources should be directed to and will be fulfilled by the lead contact, Jessica Quintin.

### Technical contact

Questions about the technical specifics of performing the protocol should be directed to and will be answered by the technical contact, Sarah S. W. Wong.

### Materials availability

This protocol does not generate new unique reagents.

### Data and code availability

This protocol does not generate or analyse any datasets or codes.

## ACKNOWLEDGMENTS

We thank J. Iñaki Guijarro from the Biological NMR and HDX-MS Technological Platform, Institut Pasteur, for support with NMR and data analysis. We also thank the Central Animal Facility of Institut Pasteur for their assistance with mouse housing.

This work was supported by the Agence Nationale de la Recherche (ANR-16-CE15-0014-01), EU Horizon 2020 research and innovation program (847507), Sanofi iAWARDS EU 2023-2024, Coordenação de Aperfeiçoamento de Pessoal de Nível Superior (CAPES, finance code 001), and Institut Pasteur (Institut Carnot–Microbes et Santé grant n°11 CARN-017-01, DARRI/DM INNOV 192-23 and PTR 617-23).

For the purpose of open access, the author has applied a CC-BY public copyright license to any Author Manuscript version arising from this submission.

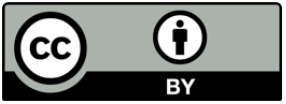

## Author contributions

Conceptualization: K.M., S.S.W.W., M.B., A.H., and J.Q.; Methodology: K.M., S.S.W.W., M.B., A.H., and J.Q.; Formal analysis: K.M., S.S.W.W., M.B., A.H., and J.Q.; Investigation: K.M., S.S.W.W., M.B., A.H., and J.Q.; Resources: J.Q.; Writing – original draft: K.M., S.S.W.W., and J.Q.; Writing – review & editing: K.M., S.S.W.W., M.B., A.H., and J.Q.; Visualization: K.M. and J.Q.; Supervision: J.Q.; Project administration: J.Q.; Funding acquisition: J.Q.

